## Supplementary figures and images for "Inference of Gene Regulatory Network Uncovers the Linkage Between Circadian Clock and Crassulacean Acid Metabolism in *Kalanchoë fedtschenkoi*"

### Supplemental Figure 1

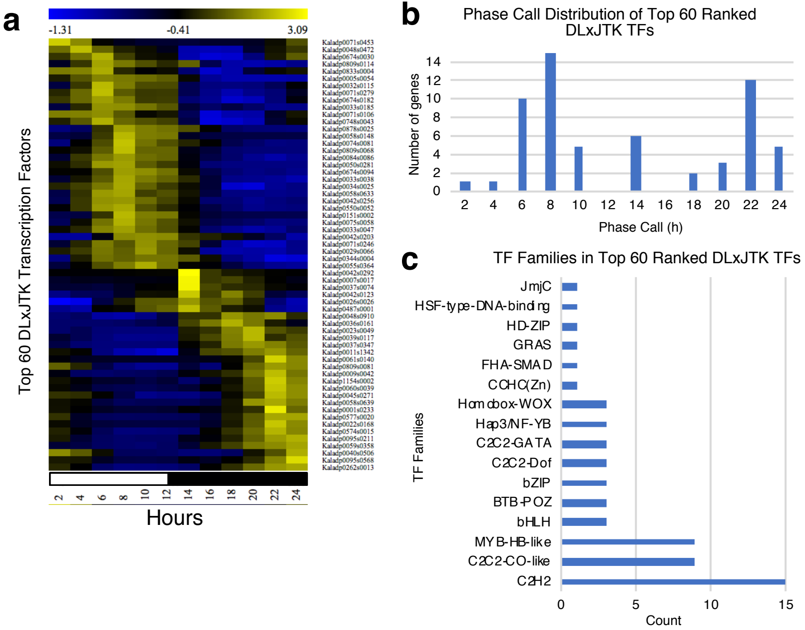
